# Rapid and efficient optical tissue clearing for volumetric imaging of the intact and injured spinal cord in mice

**DOI:** 10.1101/2025.03.22.644730

**Authors:** Andrew W. Buxton, Frank L. Jalufka, Margaret E. Hruska, Jack R. Kubaney, Dylan A. McCreedy

## Abstract

Tissue clearing and 3D imaging have emerged as powerful techniques to assess the cellular and tissue-level architecture of the spinal cord. With the rapidly increasing variety and complexity of optical tissue clearing techniques, there is a critical need for optimization and streamlining of tissue-specific protocols, particularly when dealing with injury or disease states. We evaluated and combined multiple organic solvent-based techniques to develop sciDISCO: a spinal cord injury-optimized DISCO tissue clearing protocol. sciDISCO allows for the robust clearing, labeling, and 3D imaging of the intact spinal cord, as well as clearing around and through the lesion site formed after contusive spinal cord injury. In addition, we have identified alternatives for hazardous chemicals commonly used in organic solvent-based clearing including dichloromethane and dibenzyl ether. In this study, we demonstrate the compatibility of sciDISCO with multiple different labeling techniques to provide robust analysis of unique neuronal populations and morphologies in addition to cellular and tissue-level changes occurring following spinal cord injury.

## INTRODUCTION

As the critical mediator of sensory and motor communication between the body and the brain, the spinal cord is a high priority area for neurological research. The spinal cord is composed of many phenotypically and morphologically distinct neuronal and glial populations as well as axon tracts from supraspinal, propriospinal, and peripheral neurons^1–3^. Disease and mechanical injury to spinal neuron populations or the myelinated axon tracts can disrupt signals coming from or going to the brain, leading to motor and sensory dysfunctions^4,5^. Understanding how spinal neuron populations transmit and modify motor, sensory, and autonomic signals in the intact spinal cord, as well as the cellular and circuit changes that occur in disease and injury, is critical for the development of new treatments for individuals with SCI^6–8^.

Historically, study of neuronal populations and tracts within the spinal cord has been achieved using mechanical sectioning techniques utilizing bladed instruments such as vibratomes or cryostats. While the data collected from physical tissue sections has proven highly valuable in our understanding of spinal cord circuits and function, the process can be prone to tissue deformations and is very labor-intensive. Recent technological advancements have been made to section and image spinal cord tissue in a more efficient manner^9^, but are still prone to the limitations of mechanical sectioning including tearing or distortions of tissue sections^10,11^. In contrast with the historical sectioning techniques, the emergence of optical tissue clearing has provided a fast and efficient method of imaging whole, intact tissues using virtual or optical tissue sections that can then be rapidly assembled back into a 3D image^12–17^. With tissue clearing, cells and tissue components can be visualized in their native 3D configuration with minimal tissue loss^18,19^. Over the past decade, many different forms of optical tissue clearing protocols have emerged.

Tissue clearing protocols have been commonly categorized based on their mechanism of action or properties of the solvents utilized. Categories of tissue clearing protocols include the aqueous-based methods, such as CLARITY^20^,CUBIC ^21^, and PACT^22–24^, which utilize aqueous detergents to delipidate tissues. The other major category of tissue clearing protocols are hydrophobic or organic solvent-based methods, such as 3DISCO^25,26^, iDISCO+^27,28^, uDISCO^29^, fDISCO^30^, Ethanol-ECi^31,32^, and PEGASOS^33^, which utilize organic solvents to delipidate and match the refractive indices of tissues. Collectively, the various tissue clearing protocols provide an essential tool for the 3D imaging of many different tissues. However, the abundance of protocols and inherent differences between each method often results in the need for further optimization to achieve successful tissue clearing.

In this study, we sought to develop an optical tissue clearing protocol specifically designed for studying the intact and injured spinal cord. We aimed to address several limitations commonly encountered with optical clearing of the spinal cord including the need to remove the meninges due to tissue swelling, poor clearing of the extracellular matrix (ECM)-rich lesion site following contusive spinal cord injury, curvature of the spinal cord that hinders efficient imaging, lengthy protocol times, and use of hazardous chemicals. By combining multiple organic solvent-based methods, including iDISCO+ and Ethanol-ECi, we developed the sciDISCO (<u>s</u>pinal <u>c</u>ord injury-optimized DISCO). The sciDISCO tissue clearing protocol has a shorter tissue processing time than iDISCO+ and other aqueous-based protocols, allows for the immunolabeling and clearing of the meningeal layer around the spinal cord, clears through the dense ECM that forms in the lesion site after contusive spinal cord injury (SCI), straightens the spinal cord to reduce imaging time and data size, replaces dichloromethane and dibenzyl ether with less hazardous alternatives, and allows for 3D visualization and imaging of the entire spinal cord using lightsheet microscopy. We demonstrate the utility of sciDISCO for visualizing lesion formation and neuronal loss, quantification of immune cell accumulation, imaging of the lymphatic system in the meninges of the spinal cord, and morphological analysis of sparsely-labeled neuronal populations. We believe that sciDISCO provides a robust and time efficient method for optically clearing the spinal cord that will accelerate research on normal spinal cord function as well as SCI and other spinal cord afflictions.

## Materials and Equipment

### Delipidation

1. 5 mL conicals (Axygen^©^, SCT-5ML)
2. Methanol (Fisher Chemicals, A452-1)
3. Dichloromethane (DCM, AcroSeal^©^, 326850010)

- Alternative: Benzotrifluoride (BTF, Thermo Scientific™, AAB2134030)
4. 10X phosphate-buffered saline (Corning^®^, 20-031-CV)
5. Nutating rocker — LabDoctor™ Mini Nutating Rocker (Ward’s Science™, H3D1020)
6. Tween-20 (Thermo Scientific™, J20605-AP)
7. Heparin (Sigma-Aldrich^Ⓡ^, H3393-100KU)
8. Sodium Azide (Millipore Sigma, S2002)

### Immunofluorescent Labeling

1. Normal Donkey Serum (Millipore Sigma, D9663)
2. Dimethyl Sulfoxide (Fischer Bioreagents, BP231-1)
3. 0.45 μm Syringe Filter Unit (Millex-HP, SLHPR33RS)
4. 10 mL syringe (Fisherbrand™, 14955458)
5. 1.5 mL microcentrifuge tubes (Denville, C2170)
6. 15 mL conicals (Falcon™, 352196)
7. Incubator — Ward’s® Mini Digital Incubator (Ward’s Science™, 470230-608)
8. Primary antibodies —Rabbit anti-NeuN (1:500, Millipore Sigma, ABN78), Chicken anti-NeuN (1:500, Millipore Sigma, ABN91), Rabbit anti-Lyve1 (1:500, Abcam, ab14917), Goat anti-mCherry/RFP (1:500, SicGen, AB0040-200), Goat anti-S100a8/MRP8 (1:1000, R&D Systems, AF3059) Rabbit anti-V5 (1:100, Bethyl Laboratories, A190-120A)
9. Secondary antibodies (1:200) — Donkey anti-Rabbit IgG AlexaFluor® 488 (Invitrogen, A21206), Donkey anti-Rabbit IgG AlexaFluor® 647 (Invitrogen, A32795), Donkey anti-Chicken IgG AlexaFluor® 488 (Jackson, 703-545-155), Donkey anti-Goat IgG AlexaFluor® 647 (Invitrogen, A32849),

### Agarose Mounting and Refractive Index Matching

1. Sodium phosphate, monobasic, monohydrate (Millipore Sigma, S9638)
2. Sodium phosphate, dibasic, anhydrous (Millipore Sigma, 567550)
3. Agarose, low-melting temperature (Millipore Sigma, A9045)
4. Heat block with 1.5 mL adaptor
5. Ethyl Cinnamate (Sigma-Aldrich, 112372-100G)

- Alternative for Dibenzyl Ether

## METHODS

### Animals

Adult male and female C57Bl/6N (Charles Rivers), Chx10-Cre (obtained from Dr. Steven Crone), Confetti (The Jackson Laboratory, Gt(ROSA)26Sor^tm1(CAG-Brainbow2.1)Cle^)^34^ and MORF3 (The Jackson Laboratory, Gt(ROSA)26Sor^tm3(CAG-smfp-V5*)Xwy^)^35,36^ were used for this study. Chx10-Cre were maintained on a C57Bl/6J background and bred with Confetti or MORF3 mice to obtain sparse labeling of V2a interneurons. All studies were approved by the Institutional Animal Care and Use Committee at Texas A&M University.

### Spinal Cord Injury

Adult male and female mice were anesthetized with 2% isoflurane before a laminectomy was performed at the T9 vertebrae level to expose the thoracic spinal cord. A 60 kDyne contusion injury was delivered to the spinal cord with a 1 sec dwell time using the Infinite Horizons impactor (Precision Systems and Instrumentation). Surrounding musculature was sutured closed and the skin was closed with wound clips. Manual bladder expression and antibiotic/saline subcutaneous injections were performed twice daily until the spinal cords were collected.

### Transcardial Perfusion and Fixation

Transcardial perfusion was performed as described previously^23^. Mice are deeply anesthetized with a lethal dose of 2.5% Avertin (0.02 mL/g body weight; Sigma) before being perfused with 25 mL of ice-cold 1X phosphate buffered saline (PBS). Following the perfusion with PBS, animals were then perfused with 25 mL of ice-cold 4% paraformaldehyde in PBS for fixation. After fixation, the spinal column was removed and placed in ice-cold 4% paraformaldehyde overnight for post-fixing. Following post-fixation, the spinal column was washed with 1X PBS. Using fine tweezers and microscissors, the spinal cord was carefully removed from the spinal column as previously described^23^ and was stored at 4°C in PBS and 0.01% (w/v) sodium azide until ready for clearing.

### 3D Printing of Spinal Cord Straightener and Agarose Mounting Chamber

The spinal cord straightener device and strap were 3D printed using natural polypropylene filament (Recreus PP3D) on a Prusa MK3S+ printer with a layer height of 0.10 mm. Polypropylene filament was chosen due to its chemical resistance to the organic solvent clearing solutions. Packing tape was placed on the printer bed to increase adhesion of the polypropylene filament during printing. The agarose mounting chamber was 3D printed using polylactic acid (PLA) filament (Amazon Basics PLA) on a Prusa MK3S+ with a layer height of 0.10 mm.

### Passive CLARITY and iDISCO+ Tissue Clearing

To compare our tissue clearing protocol to current aqueous and organic based clearing protocols, we performed PACT and iDISCO+ tissue clearing as previously described^22–24,28,37^ on spinal cords from adult male and female wild-type (WT) mice.

### sciDISCO Tissue Clearing

#### Delipidation

##### Purpose

The delipidation step removes light scattering lipids from the tissue.

1. Dehydration of the spinal cord

1.1. Transfer the dissected spinal cord from the 1X PBS solution into the 3D-printed spinal cord polypropylene straightener and secure it with the 3D-printed polypropylene strap. The spinal cord will remain in the straightener throughout the delipidation and rehydration process.
1.2. Prepare the dehydration series in 5 mL conicals of methanol and ddH_2_O: 20%, 40%, 60%, 80%, and 100% MeOH/ddH_2_O (v/v).
1.3. Transfer the spinal cord to the 20% MeOH/ddH_2_O solution, cap the conical tightly to prevent leaks, and rock on a nutating rocker for 1 hour at room temperature. After the hour, transfer to the 40% MeOH/ddH_2_O solution and rock for another hour. Repeat throughout the entire dehydration series, rocking the spinal cord in each solution for 1 hour at room temperature. After the final 100% MeOH/ddH_2_O solution, transfer the spinal cord to a new conical containing 5 mL of 100% MeOH and rock for 1 hour at room temperature to ensure complete removal of water from the sample.
1.4. Prepare 5 mL of 66% DCM/ 33% MeOH (v/v) solution in a 5 mL conical. BTF can be used instead of DCM. Use glass stripettes and polypropylene tubes when working with DCM or BTF.
1.5. Transfer the spinal cord to the prepared DCM/MeOH solution and incubate overnight while gently rocking on the nutating rocker at room temperature. BTF can be used instead of DCM.
2. Rehydration of the spinal cord

2.1. Transfer the spinal cord from the 66% DCM/ 33% MeOH solution to a 5 mL conical filled with 100% methanol. Incubate the spinal cord in 100% MeOH for 30 minutes with gentle rocking, then replace the MeOH with 5 mL of fresh 100% MeOH and incubate for an additional 30 minutes with gentle rocking at room temperature.
2.2. Prepare the rehydration series (MeOH/H_2_O) in 5 mL conicals with the following percentages of methanol in ddH_2_O: 80%, 60%, 40%, and 20% MeOH/ddH_2_O (v/v).
2.3. Rehydrate the spinal cord sample by placing the spinal cord in the rehydration series starting with the 80% MeOH/ddH_2_O solution for 1 hour rocking on a nutating rocker at room temperature. After the hour, transfer the tissue through the remainder of the series, 1 hour in each solution, rocking on a nutating rocker at room temperature.
2.4. Prepare 1x PBS.

2.4.1. Combine 100 mL of 10X PBS with 900 mL of ddH_2_O and stir.
2.5. Transfer 5 mL of the prepared PBS solution to the 5 mL conical. Place the spinal cord sample in this PBS solution for 30 minutes with gentle rocking at room temperature.
2.6. Prepare 1 L of PBS/Tween-20 with Heparin (PTwH)

2.6.1. Combine 100 mL of 10X PBS, 2 mL Tween-20, and 1 mL of 10 mg/mL Heparin and ddH_2_O to a final volume of 1 L. Stir until thoroughly mixed. Add sodium azide to a final concentration of 0.02% (w/v). Store at room temperature.
2.7. Remove the spinal cord from the straightener and transfer 5 mL of the prepared PTwH into a 5 mL conicals. Wash the sample in PTwH for 30 minutes. Repeat.

###### Checkpoint

Possible stopping point – tissue may be stored in 1x PBS with 0.01% (w/v) sodium azide at 4°C for several weeks.

#### Immunofluorescent Labeling

##### Purpose

Immunofluorescent labeling allows for the visualization of endogenous or transgenic

epitopes within the cleared tissue.

1. Primary antibody staining

1.1. Prepare 1 mL of the primary antibody solution in a 1.7 mL microcentrifuge tube.

1.1.1. Combine: PTwH, 5% DMSO, 3% Normal Donkey Serum, and primary antibodies. If samples are too large for 1.5 mL microcentrifuge tubes, use a 5ml conical and adjust volume of primary antibody solution accordingly.
1.2. Incubate the spinal cord sample in the primary antibody solution for 2 days at 37°C with gentle rocking.
2. PTwH washes

2.1. Wash the spinal cord sample 5 times for 30 minutes each with 12.5 mL of PTwH in a 15 mL conical while rocking with a nutating rocker at room temperature.
2.2. Leave the sample in the 5th wash overnight with gentle rocking at room temperatue.
3. Secondary antibody staining

3.1. Prepare 1.2 mL of the secondary antibody solution in a 1.7 mL microcentrifuge tube.

3.1.1. Combine PTwH, 5% NDS, and secondary antibodies.
3.1.2. Once prepared, filter the secondary antibody solution using a 3 mL syringe attached to a 0.22 µm syringe filter. Filter into a new 1.7 mL microcentrifuge. Some loss of the secondary antibody solution will occur, but the final volume should be ∼ 1 mL.
3.2. Incubate spinal cord samples for 2 days at 37°C in the secondary antibody solution with gentle rocking.
4. Wash samples in PTwH

4.1. Repeat PTwH Washes (Step 2), leaving the sample in the 5th PTwH wash overnight on the nutating rocker at room temperature.

#### Agarose Mounting and Refractive Index Matching

##### Purpose

Mounting spinal cord samples in agarose can help with positioning and manipulating the sample safely within the Zeiss lightsheet microscope. Refractive index matching helps homogenize the refractive indices of the tissue and agarose to minimize light diffraction during imaging.

1. Mounting spinal cord in agarose (optional)

1.1. Prepare 0.1 M phosphate buffer (PB), pH 7.4.

1.1.1. Dissolve 3.1 g of sodium phosphate (NaH_2_PO_4_, monobasic, monohydrate) and 10.9 g of sodium phosphate (Na_2_HPO_4_, dibasic, anhydrous) in 900 mL of ddH_2_O. Add ddH_2_O to a final volume of 1L.
1.2. Prepare 0.02 M phosphate buffer by combining 200 mL of 0.1 M PB with 800 mL of ddH_2_O.
1.3. Prepare sufficient volume of 2% agarose to fully immerse the spinal cord and any microscope attachments necessary for moving the sample.

1.3.1. To make 20 mL of 2% agarose, combine 400 mg of low-melt agarose in 20 mL of 0.02 M phosphate buffer.
1.3.2. Microwave in increments to avoid boiling over until all the agarose is fully dissolved.
1.3.3. Aliquot 1 mL into 1.5 mL microcentrifuge tubes. Aliquots can be stored at 4°C.
1.3.4. To re-melt, place microcentrifuge tubes in the heat block at 95°C. Cool the agarose to ∼60°C prior to use.
1.4. Position the spinal cord and any microscope handling components in a mold for the agarose. Slowly pipette in melted agarose until the sample and components are fully submerged. Allow agarose to solidify in a light protected environment for at least 30 minutes. Do not let agarose set for more than 2 hours as it can dry out and crack. Once sample is embedded, proceed to the next step.
2. Dehydration of the spinal cord

2.1. Prepare the dehydration series (MeOH/H_2_O) in 5 mL conicals with the following concentrations of methanol: 20%, 40%, 60%, 80%, and 100%.
2.2. Dehydrate the sample through the prepared series in increasing order of methanol concentration.

2.2.1. Lay the conicals horizontally on a nutating rocker to prevent the spinal cords and agarose from warping along the bottom of the conicals during the dehydration steps. Gently rock for 1 hour each at room temperature.
2.3. Incubate the spinal cord sample in 100% MeOH overnight in a 5 mL conical with gentle rocking at room temperature.
3. Refractive index matching

3.1. Prepare 5 mL of 66% DCM/ 33% MeOH (v/v) solution in a 5 mL conical. BTF can be used instead of DCM.
3.2. Incubate sample in the DCM/MeOH solution for 3 hours on a nutating rocker at room temperature.
3.3. Wash sample twice in 5 mL of 100% DCM in 5 mL conicals for 15 minutes each on the nutating rocker at room temperature. BTF can be used instead of DCM.
3.4. Prepare 5 mL of ethyl cinnamate in the 5 mL conical.
3.5. Transfer the spinal cord sample to the ethyl cinnamate solution and equilibrate for at room temperature for at least 1 full day prior to imaging. For best results, gently invert the conical every 1-2 hours before leaving overnight.

3.5.1. The following day, replace with fresh ethyl cinnamate and let the sample rest for at least 2 hours prior to imaging. Gently invert the conical every 1-2 hours to ensure mixing of the ethyl cinnamate throughout the sample.

### Lightsheet Microscopy

After equilibration in ethyl cinnamate for at least 1 day, spinal cord samples were imaged using a Zeiss Z1 Lightsheet microscope with a 5x acquisition objective. Virtual image z-stacks were captured using both right and left sided lightsheet illumination (5x illumination objectives) and fused together in Zen software.

### Image Analysis

Lightsheet images were analyzed in Imaris 3D software (Bitplane), Matlab, Python, or ImageJ^38^. To calculate the mean fluorescent intensity of NeuN labeling, an image mask was created for NeuN^+^ pixels for each image in MATLAB. The NeuN^+^ image mask was used to generate a binary image file removing any background fluorescence. The masked image file was then imported into a Python script, which was used to calculate the average fluorescent intensity of each image. To calculate the signal-to-background ratio of NeuN labeling, a signal mask for the NeuN^+^ pixels was created using a manual threshold, and a background mask was generated by thresholding the entire tissue and subtracting the signal mask. The same signal and background threshold levels were used for every image. The signal-to-background ratio for each image was calculated by dividing the mean intensity within the signal mask by the mean intensity within the background mask. MRP8^+^ cell quantification was performed by using the Imaris spot function. The number of MRP8^+^ cells within 500 µm of the lesion center were counted in each image.

### Statistical Analysis

Statistical analysis was performed using Prism software (Graphpad). Sample sizes for replicates are presented in figure legends. Data is presented as mean ± SEM. P values were calculated by two-sided Student’s *t* test and one-way ANOVA with Tukey’s post hoc test. P < 0.05 was considered significant.

## RESULTS

### sciDISCO clears the full intact murine spinal cord in a time efficient manner

To develop a streamlined tissue clearing protocol, we identified and combined the essential and most effective components of multiple tissue clearing procedures including iDISCO+ and ethanol-ECi^26,28,39^. The resulting sciDISCO optical clearing protocol is comprised of three main steps: delipidation, tissue labeling, and refractive index matching (Fig. 1A). The first step involves delipidation, in which methanol and dichloromethane (DCM) or benzotrifluoride (BTF) are used to remove the majority of light scattering lipids from the tissue. Next in the immunolabeling step, primary and secondary antibodies are used to label target epitopes in order visualize biomolecules with cellular and subcellular resolution. Finally in the refractive index matching step, the spinal cord is encased in agarose using our 3D printed agarose mounting chamber (Fig. 1B), dehydrated again, and then incubated in ethyl cinnamate for refractive index matching for at least 24 hours prior to imaging (Fig. 1D). Samples can be kept light protected at room temperature in ethyl cinnamate until imaging.

**Figure 1.**
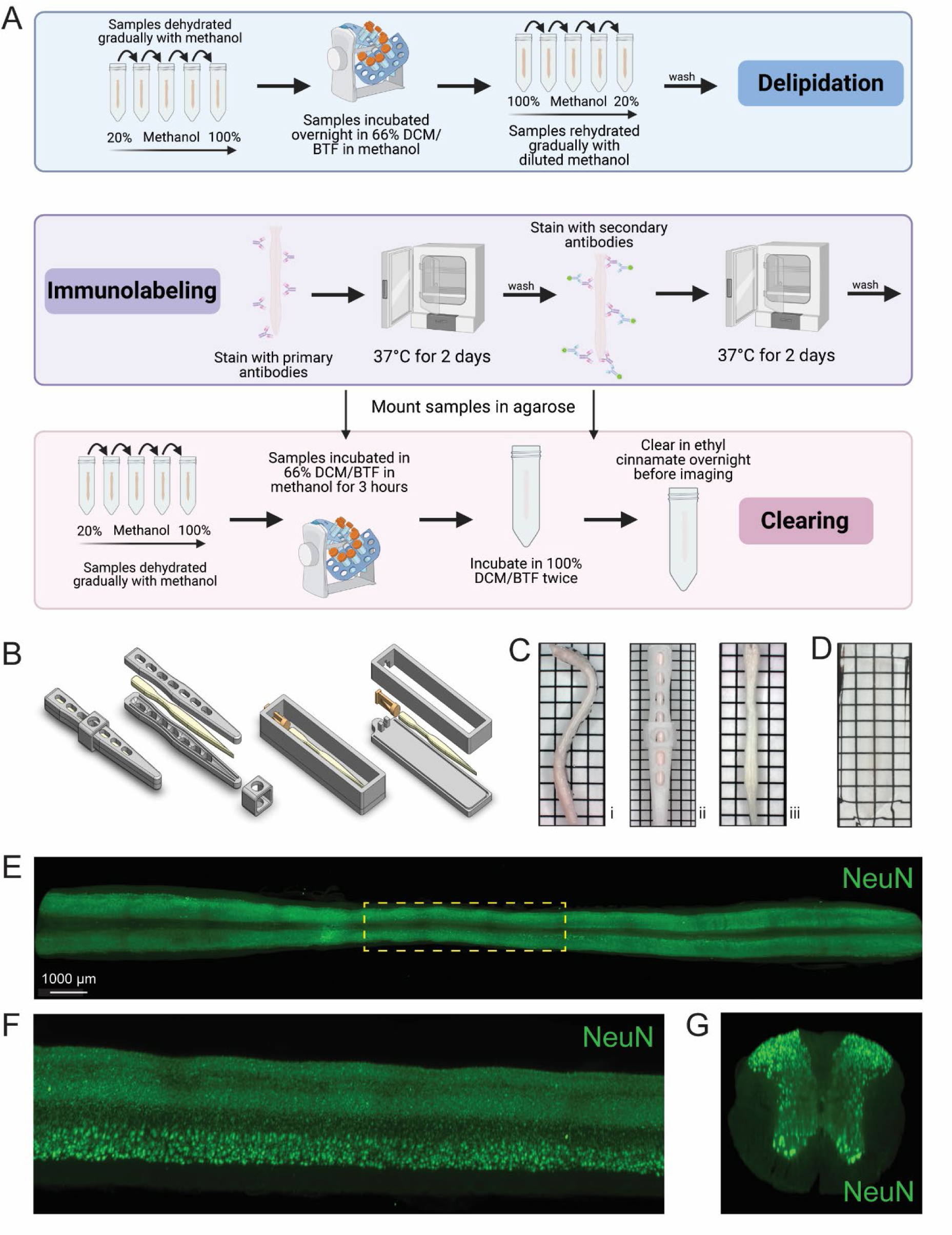
The sciDISCO protocol allows for the 3D imaging of the entire spinal cord. (A) Visual schematic of the sciDISCO protocol involving the delipidation of the spinal cord, immunolabeling, and RI matching in ethyl cinnamate. Created in BioRender. https://BioRender.com/7kqz2km. (B) 3D schematics of the spinal cord straightener and agarose mounting chamber. (C) (i) A curved spinal cord after post-fixation and dissection. (ii) The spinal cord in the 3D printed spinal cord straightener and (iii) the straightened spinal cord after delipidation and removal from the straightener. (D) A cleared spinal cord embedded in agarose and RI matched with ethyl cinnamate. (E) Horizontal view of a 3D lightsheet image of a spinal cord including cervical, thoracic, and lumbar segments labeled with NeuN antibody to visualize neuronal cell bodies. (F) Zoomed in sagittal view of the outline thoracic region (yellow) from E. (G) Virtual transverse cross section of the thoracic spinal cord. Grids on images are made up of 2.5 mm x 2.5 mm squares.

The murine spinal column is inherently curved and we noticed that the dissected spinal cords reverted to a pronounced curved shape during the dehydration step (Fig. 1Ci). Imaging the curved spinal cord is technically challenging and increases imaging time and image data size by requiring imaging of void space below and above the curved spinal cord regions. To address these limitations, we developed a 3D printed straightener device to house the spinal cord during the initial delipidation steps (Fig. 1B, 1Cii). Following delipidation using this cage, the spinal cord remained straight for the duration of the protocol (Fig. 1Ciii).

Using the sciDISCO protocol, we were able to capture immunofluorescent staining (NeuN) in the full length, intact murine spinal cord using lightsheet microscopy (Fig. 1E). The resulting 3D images enabled visualization of the entire labeled spinal cord tissue (Fig. 1E, horizontal view). Multiple view angles of selected regions of the 3D image can be readily obtained including the sagittal (Fig. 1F) and transverse views (Fig. 1G).

In respect to comparable protocols, iDISCO+ and PACT, sciDISCO is considerably more time efficient, able to clear and immunolabel the whole spinal cord in as little as 9 days, compared to 12 days for iDISCO+ and 16 days for PACT (Fig. 2A). Despite the reduced time required for clearing and immunolabeling, sciDISCO delivers comparable results using intact spinal cords with the meninges removed (Fig. 2B).

**Figure 2.**
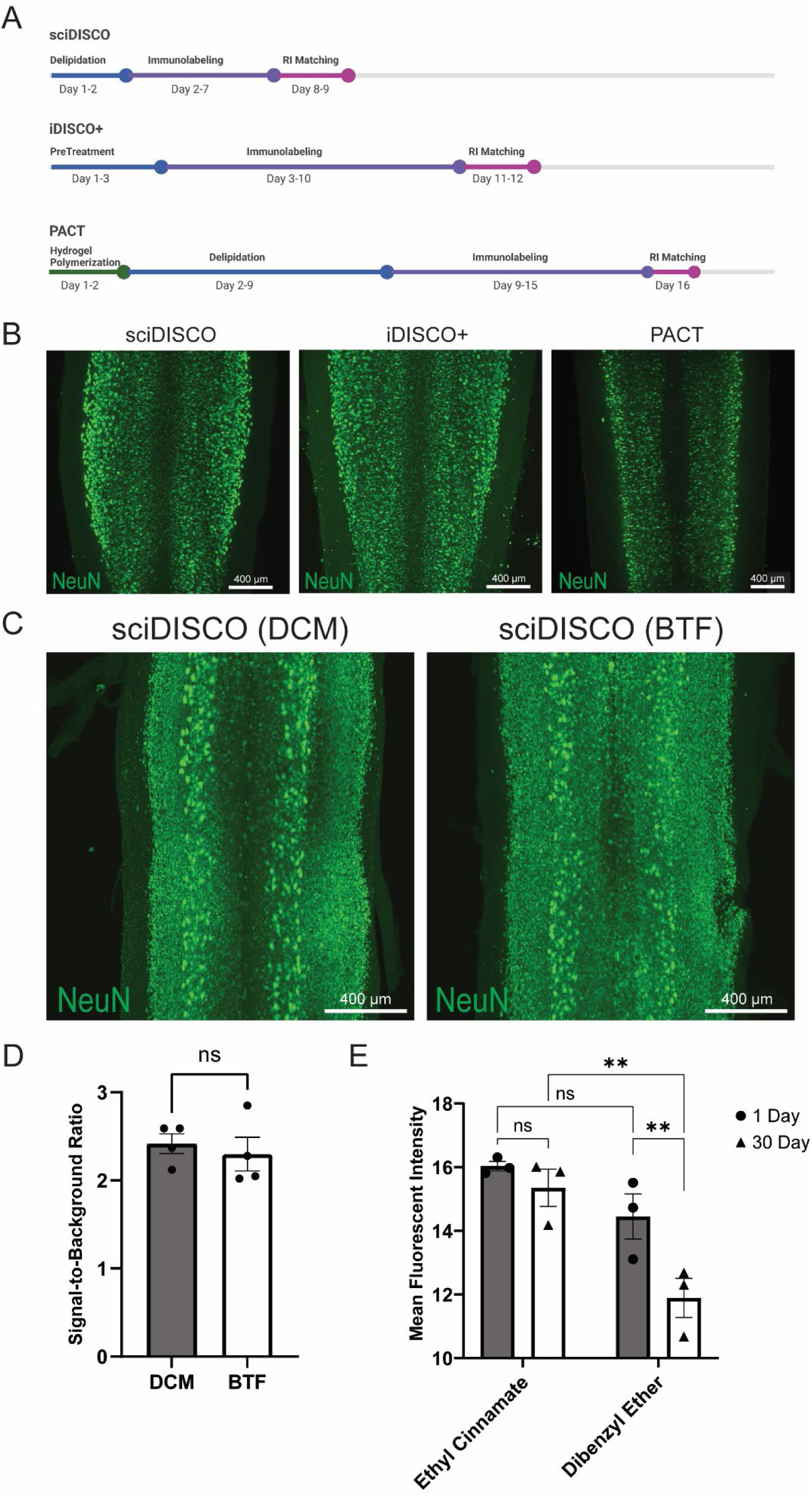
sciDISCO is faster than comparable clearing protocols and delivers similar immunolabeling. (A) Timeline schematic detailing the average times for each main step in sciDISCO, iDISCO+, and the PACT clearing protocols. Created in BioRender. https://BioRender.com/ghn1jij. (B) sciDISCO delivers similar immunolabeling of NeuN when compared to iDISCO+ and PACT clearing protocols. Representative maximum intensity projections of 3D lightsheet images of the lower cervical and upper thoracic regions are shown. (C) Representative maximum intensity projections of NeuN-labeled thoracic spinal cord regions cleared using either DCM or BTF in the sciDISCO protocol. (D) Quantification of the signal-to-background ratio of samples cleared with sciDISCO using either DCM or BTF. Mean ± SEM. Student’s t-test. (E) Quantification of mean fluorescent intensity of samples incubated in ethyl cinnamate (sciDISCO) vs dibenzyl ether (iDISCO+) after 1 and 30 days of incubation. Mean ± SEM. ** p<0.01. Two-way ANOVA with Tukey’s multiple comparisons post-hoc test.

DCM was originally used for delipidation in the development of the sciDISCO protocol. However, there are potential health and environmental risks when working with DCM^40^, as well as an upcoming ban in the United States for most uses of DCM. We found that another, more environmentally friendly organic solvent, benzotrifluoride (BTF), performs comparably when used as a direct alternative to DCM in the sciDISCO protocol (Fig. 2C). BTF also maintains the same signal-to-background ratio of NeuN labeling compared to DCM with sciDISCO (Fig. 2D). We also wanted to test the long-term storage capability of samples in RI matching solutions for sciDISCO vs iDISCO+. We allowed samples to incubate in ethyl cinnamate and dibenzyl ether, respectively, in light protected boxes at room temperature for 30 days prior to imaging. Maximum intensity projections were generated from 3D images for each sample and a mask of all NeuN^+^ pixels was generated using MATLAB. The NeuN^+^ areas of each image were then assessed in Python to calculate the mean fluorescent intensity of all NeuN^+^ voxels for each sample. After 1 day of refractive index matching, there was no significant difference in mean fluorescent intensity of NeuN labeling between the sciDISCO samples stored in ethyl cinnamate and the iDISCO+ samples stored in dibenzyl ether (Fig. 2E). However, after 30 days of incubation the iDISCO+ samples had reduced fluorescent intensity relative to sciDISCO samples as well as iDISCO+ samples at 1 day of incubation. No decline in fluorescent intensity was observed with sciDISCO samples stored in ethyl cinnamate.

### sciDISCO clears through the lesion in the sub-acutely injured spinal cord

Following contusive SCI in mice, the lesion site undergoes extensive remodeling and becomes filled with ECM proteins such as collagen IV and is surrounded by a layer of reactive astrocytes commonly referred to the as the glial border or scar (Fig. 3A)^41,42^. The ability to clear through the lesion site would allow for the 3D analysis of lesion formation and structure. Previous studies have demonstrated limited clearing of the lesion site after the sub-acute phase (10-14 dpi) with aqueous-based protocols such as PACT or CLARITY^2,23^. We also observed poor clearing of the lesion site with PACT, however, the sciDISCO protocol readily cleared through the ECM-rich lesion at 14 dpi (Fig. 3B).

**Figure 3.**
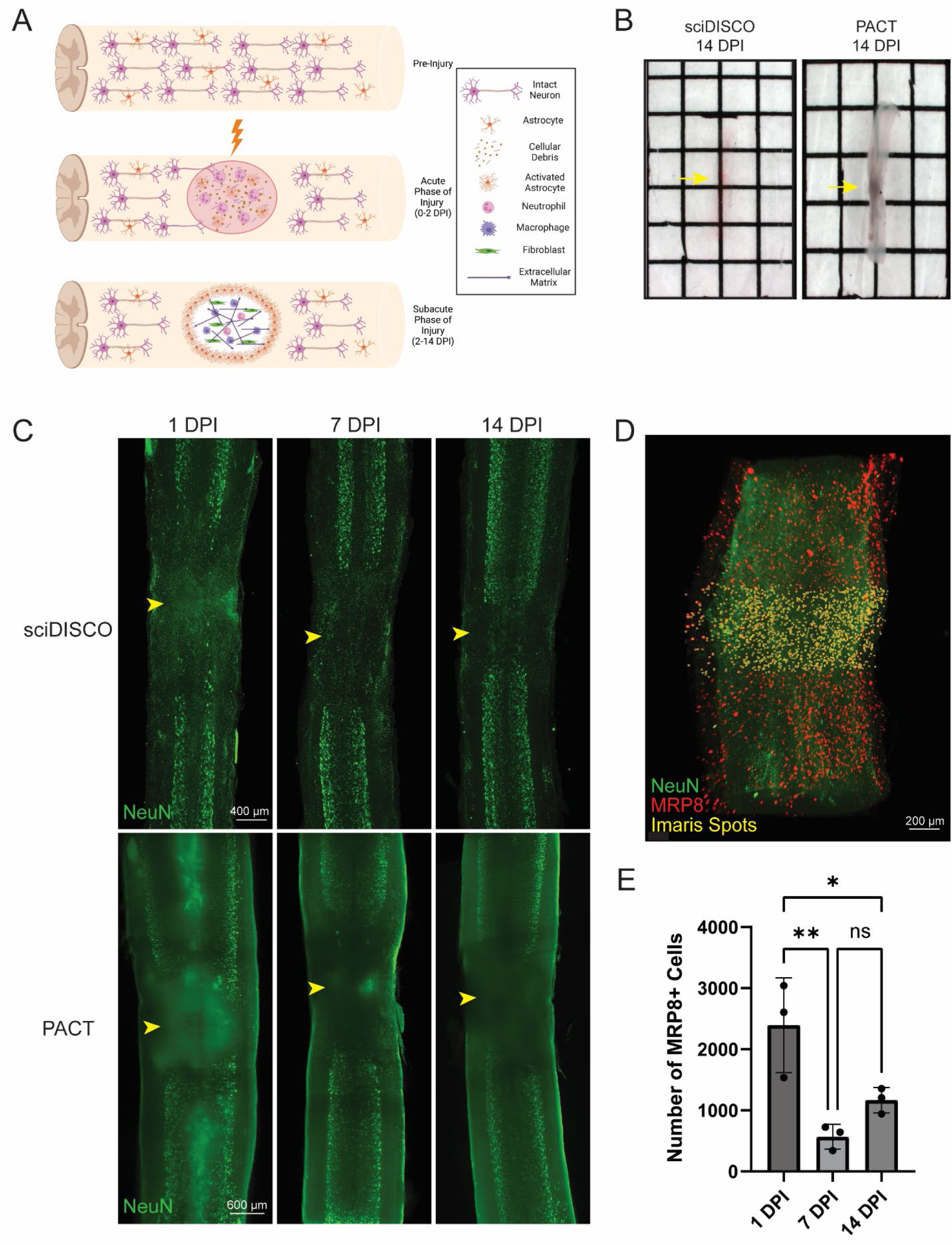
sciDISCO clears through the subacute lesion after contusive spinal cord injury. (A) Graphical representation of the lesion formation and extracellular matrix deposition following contusive spinal cord injury in mice. By the subacute phase of injury, the lesion site is filled with extracellular matrix proteins, macrophages, and fibroblasts. Created in BioRender. https://BioRender.com/yzaetus. (B) Macroscopic images of thoracic spinal cords at 14 dpi. Samples are RI matched in ethyl cinnamate (sciDISCO) or RIMS (PACT) with the lesion site indicated by the yellow arrow. The lesion site remains opaque in the PACT sample. (C) Maximum intensity projections of sciDISCO and PACT cleared thoracic spinal cords at 1, 7, and 14 days post contusive injury. The lesion center is indicated by arrowhead. (D) Example image of quantification of MRP8^+^ cells within 500 μm of the lesion center using the Imaris spots function. (E) Quantification of MRP8^+^ cell counts in sciDISCO cleared cords at 1, 7, and 14 dpi. *p<0.05, **p<0.01. One-way ANOVA with Tukey’s post-hoc test.

The increased capacity to clear through the sub-acute lesion site led to improved lightsheet imaging of the injured spinal cord cleared with sciDISCO relative to PACT (Fig. 3C). Beginning at as early as 1 dpi, the SCI lesion site failed to fully clear with PACT leading to opaque regions in the lightsheet images due to light scattering. Acutely injured spinal cord tissues readily cleared with sciDISCO, leading to virtually no opaque regions in the resulting lightsheet images. With PACT clearing, the light scattering effect of the poorly cleared lesion site is more pronounced at 7 and 14 dpi timepoint as the lesion site is remodeled and the deposited ECM is resistant to the aqueous clearing technique^13^. Despite the increased presence of ECM, sciDISCO readily cleared through the SCI lesion site of the subacutely injured spinal cord.

To test the ability to quantify cells within the lesion after clearing, we labeled neutrophils in the injured spinal cord with an anti-MRP8 antibody and quantified the number of neutrophils within 500 µm of the lesion center using Imaris (Fig. 3D-E). Neutrophils were used as our target cell as they are among the first peripheral immune cells to infiltrate in large numbers following SCI and have been shown to peak in number at 1 dpi before gradually decreasing over time in the injury site^43,44^. Using the Imaris spots function, MRP8-positive cells were quantified within the lesion site for 1, 7, and 14 dpi spinal cords. Quantitative analysis showed that there is an initial influx of neutrophils at the 1 dpi timepoint that decreases at the 7 and 14 dpi timepoints (Fig. 3E). Overall, these results indicate that our sciDISCO protocol is efficient in clearing through the acutely and sub-acutely injured spinal cord and allows for the quantification and study of inflammation and lesion formation after injury.

### sciDISCO allows for immunolabeling and clearing of the meningeal layers around the spinal cord

The meninges comprise a set of protective membranes that surround the central nervous system, including the spinal cord, and contain lymphatic vessels that play a crucial role in immune surveillance and the transport of immune cells^45^. The meningeal layers of the spinal cord have also been recently identified as a critical location for immune cell trafficking in mouse models of SCI or experimental autoimmune encephalitis^46,47^. However, many protocols require the removal of meninges to improve optical transparency and allow for swelling of the spinal cord tissue parenchyma during the tissue clearing procedures. We have previously found that the meningeal layer had to be removed from the fixed spinal cord prior to PACT clearing^23^. Without removal, the meninges frequently constricted around the swelling tissue and prevented adequate tissue clearing and antibody penetration (Fig. 4A-B). While removal of the meninges improved PACT clearing and immunolabeling, imaging of Lyve1^+^ lymphatic vessels in the meninges was severely compromised (Fig. 4B). After clearing with sciDISCO, the meninges and spinal cord cleared completely (Fig. 4C) and robust labeling of both neurons in the parenchyma and the lymphatic vessels within the meninges was readily observed (Fig. 4D). Collectively, our findings demonstrate the flexibility of sciDISCO for tissue clearing and immunolabeling with and without the meningeal layers surrounding the spinal cord.

**Figure 4.**
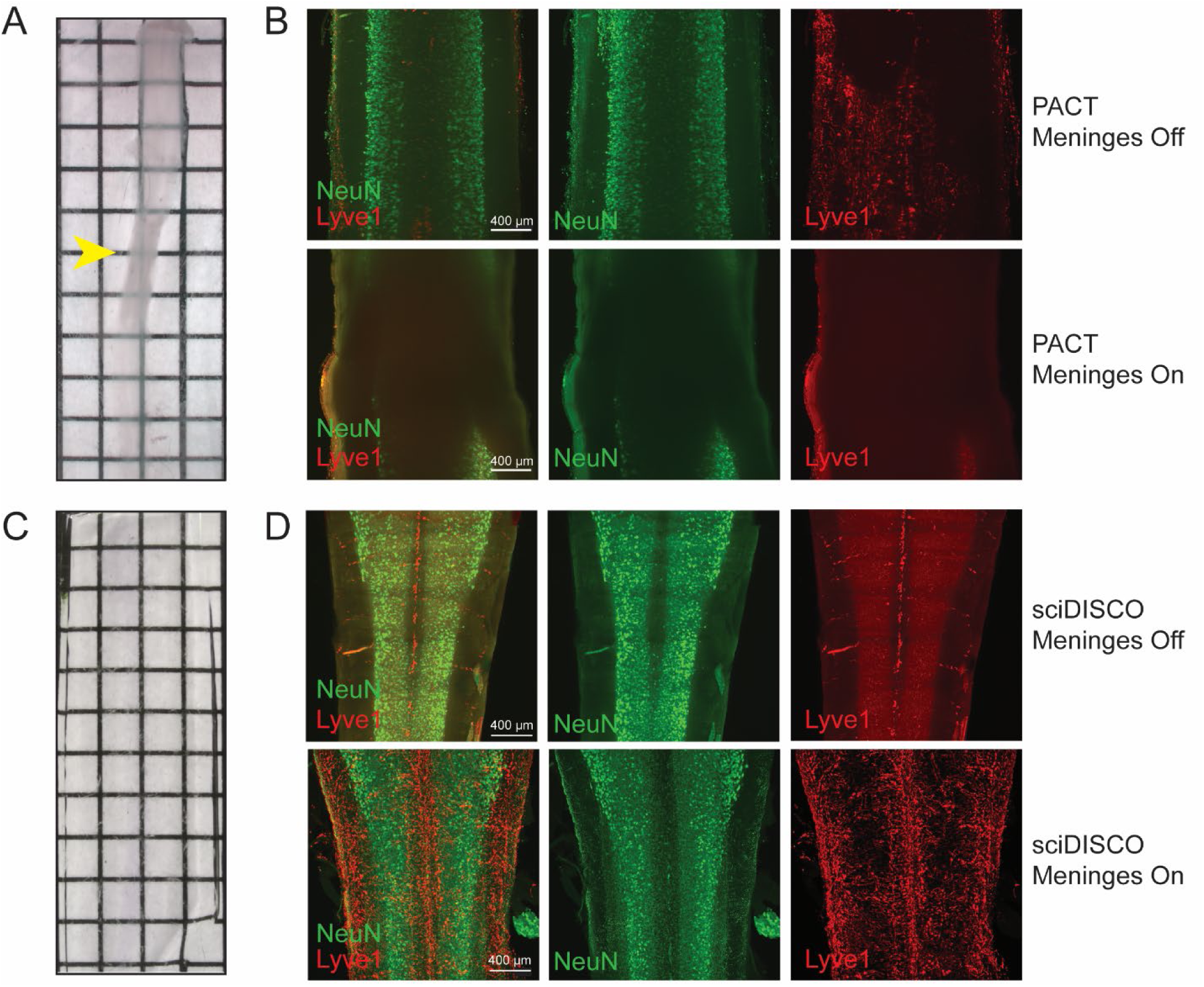
sciDISCO allows for the immunolabeling and imaging of the meningeal layers around the spinal cord. (A) Macroscopic image of a mouse spinal cord cleared and RI matched in RIMS (PACT) with the meninges still intact. The PACT clearing process causes the tissue parenchyma to swell and the meninges to constrict around the spinal cord (arrowhead). (B) NeuN and Lyve1 immunostaining of a PACT cleared spinal cord with meninges removed (top) and intact (bottom). (C) Macroscopic image of a mouse spinal cord cleared and RI matched with ethyl cinnamate (sciDISCO) with the meninges still intact. (D) NeuN and Lyve1 immunostaining of a sciDISCO cleared spinal cord with the meninges removed (top) and with the meninges intact (bottom). All images in B and D are maximum intensity projections of 3D lightsheet images. Grids on macroscopic images are made up of 2.5 mm x 2.5 mm squares.

### sciDISCO is compatible with transgenic sparse labeling techniques to identify morphological characteristics of neuronal populations

Finally, to test that our protocol is compatible with reporter mouse lines and multiple types of transgenic labels, we tested two different sparse labeling techniques to assess neuronal morphology. We first bred Chx10-Cre mice with the Confetti (*Gt(ROSA)26Sor^tm1(CAG-Brainbow2.1)Cle^*) reporter mouse line resulting in offspring (referred to as Chx10-Confetti mice) with Chx10^+^ neurons (putatively V2a interneurons) stochastically expressing either red fluorescent protein (RFP), cyan fluorescent protein (CFP), green fluorescent protein (GFP), or yellow fluorescent protein (YFP; Fig. 5A). V2a interneurons were selected as they have recently been shown to play a vital role in recovery of locomotor function after injury^8,48–51^. As with other organic solvent-based clearing protocols, endogenous fluorescence is quenched, therefore an antibody against RFP was used to label RFP-expressing V2a interneurons (Fig. 5B). This labeling strategy allowed for the visualization of V2a interneuron soma and tracts.

**Figure 5.**
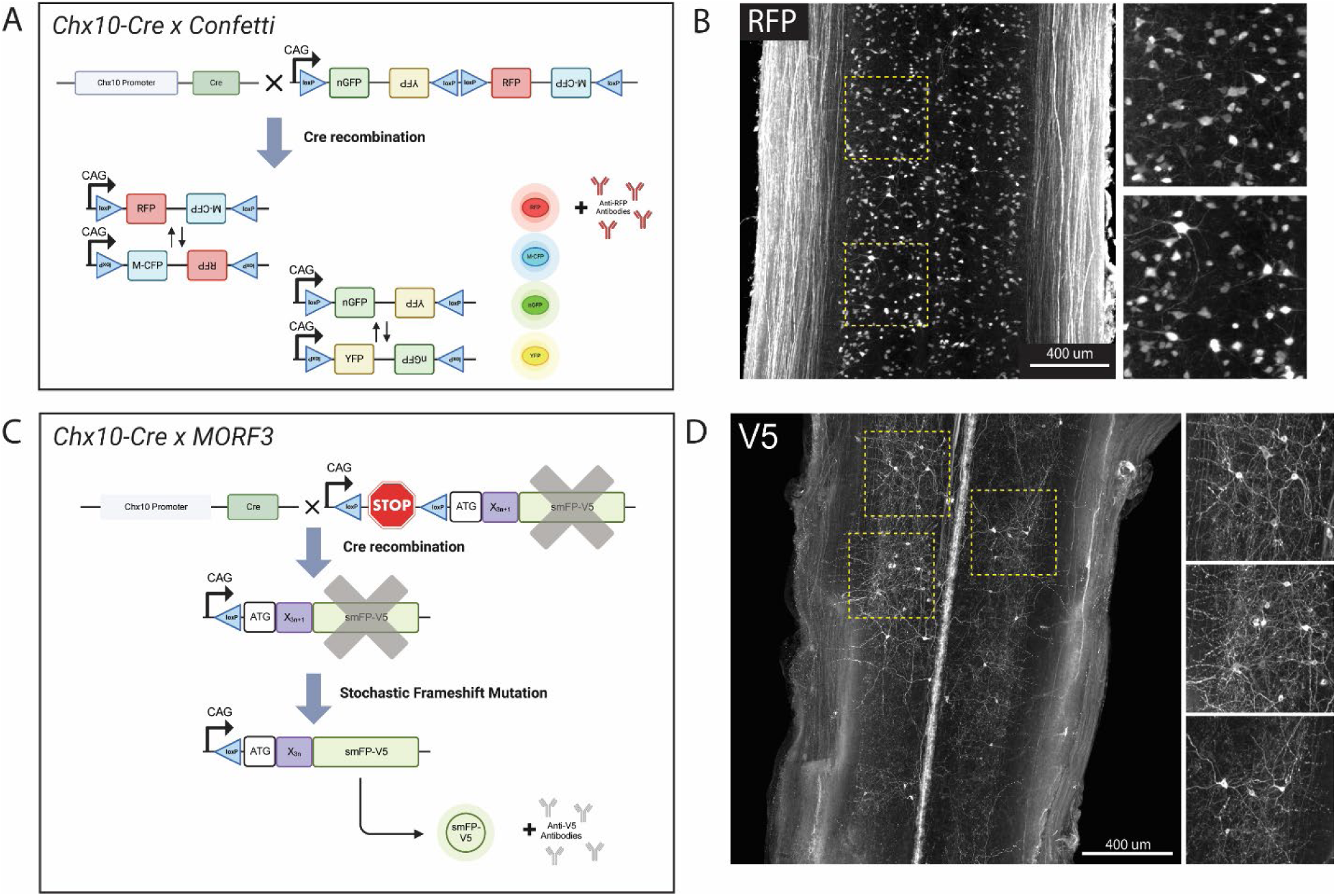
sciDISCO is compatible with transgenic mouse lines for sparse labeling of neuronal populations. (A) Graphical schematic of the stochastic Confetti fluorescent reporter expression upon crossing with Chx10-Cre mice. Created in BioRender. https://BioRender.com/5cwn62g (B) Lightsheet imaging of RFP^+^ V2a interneurons in the thoracic spinal cord following clearing with sciDISCO. (C) Graphical schematic of the MORF3 sparse labeling system upon crossing with Chx10-Cre mice. Created in BioRender. https://BioRender.com/fh674nv. (D) Lightsheet images of very sparse labeling of V2a interneurons in the thoracic region of the spinal cord following sciDISCO clearing. Both images in B and D are partial maximum intensity projections of the intermediate laminae of the spinal cord from the 3D lightsheet images.

While visualization of individual V2a interneuron soma and proximal neurites was readily accomplished with tissue clearing and lightsheet imaging of the spinal cords from Chx10-Confetti mice, long-distance tracing of axons could not be performed due to the substantial density of RFP^+^ axons in the white matter. To achieve sparser labeling of V2a interneurons, we crossed Chx10-Cre mice with MORF3 (*Gt(ROSA)26Sor^tm3(CAG-smfp-V5*)Xwy^*) reporter mice. The MORF3 mouse line contains a stop codon under control of the Cre-Lox system, and then a spaghetti monster fluorescent protein V5 (smFP-V5-F) behind a MORF (Mononucleotide Repeat Frameshift) sequence^52^. During neural development, as V2a interneurons are generated, the stop codon will be removed by Cre and a stochastic frameshift of the mononucleotide repeat region will occur in a small subset of V2a interneurons bringing the smFP-V5-F reporter protein coding sequence into frame (Fig. 5C). Immunolabeling with anti-V5 antibodies after sciDISCO clearing enabled visualization of sparsely-labeled V2a interneurons, including their cell bodies and neurites in the spinal cord (Fig. 5D). Combining sciDISCO and sparse-labeling reporter lines allows for detailed morphological analysis of spinal interneuron populations, which can be utilized for future connectomic studies.

## DISCUSSION

Optical tissue clearing has emerged as an important method for the study of 3D structures within tissues without the need for mechanical sectioning of the tissue or the requirement to rebuild a tissue from hundreds of individual tissue sections. In addition to allowing novel insights into the structure and function of normal healthy tissues, tissue clearing and 3D imaging can be highly valuable for assessing changes associated with injury or disease states. While tissue clearing techniques have been described since 1914^53^ and come in a wide range of varieties, the need for optimizations for different disease states or tissues is still warranted. By combining the most effective components of PACT, iDISCO+ and Eci-based tissue clearing procedures, we have developed sciDISCO as an optimized and streamlined protocol for optical clearing of the spinal cord, including the lesion site after SCI.

An issue that arises with respect to optical tissue clearing and lightsheet microscopy is the immense amount of image data that needs to be stored. Large-scale 3D imaging of cleared samples can generate many gigabytes to terabytes of images, particularly when dealing with larger tissues such as the spinal cord^54^. Curves or bends within the tissue can lead to excess data acquisition and storage as more void space can be required to image the entire tissue. Specifically, when dehydrating the murine spinal cord, the curvature of the spinal cord is exacerbated, which can lead to an increase in image data when imaging the entire cord. The use of our 3D printed spinal cord straightener keeps the spinal cord straight during dehydration, allowing for images to be captured with fewer imaging planes (optical sections) to image the entire cord and reducing the amount of data and time for each image, as well as lowering the required storage space.

One major area of SCI research involves the lesion and glial border that forms following traumatic SCI. In mice, the glial border forms around the lesion center in the sub-acute phase of injury. Within the lesion, a dense network of collagen fibers and other ECM proteins fills the lesion^41^, and is surrounded by border-forming astrocytes^55^. sciDISCO, with the ability to clear through this dense lesion and visualize components within the lesion and within the glial border, opens up areas of research to further understand lesion formation and glial border development over time, and understanding how the lesion and glial border interact with spared neurons around the injury site.

Following the disruption of neural circuitry caused by SCI, studies have shown that some damaged neurons display a degree of plasticity and form detour circuits, circumventing the injury and lesion area in an attempt to contribute to functional recovery^56^. Detour circuits can form from damaged neurons sprouting collaterals that establish new connections with neurons spared after the injury^57,58^. Previous research has shown long-distance propriospinal interneurons contribute to detour circuit formation and locomotor functional recovery following cervical injury^48^. sciDISCO, with the ability to clear through the lesion site, and combined with sparse neuronal labeling technologies such as the MORF3 mouse line, can allow for the visualization and mapping of detour circuit formation following SCI. In addition to detour circuit mapping, identification of morphological changes in spared neurons following SCI can lead to a better understanding of endogenous recovery mechanisms and spontaneous functional recovery within the spinal cord following injury.

The detailed protocol for sciDISCO described in this report will provide a powerful tool for studying the spinal cord, including SCI and recovery. By streamlining the procedures, we aim to make tissue clearing of the spinal cord more accessible and efficient. While lightsheet microscopy was used in this study to generate 3D images, the optically cleared spinal cord could be imaged with other modalities including confocal microscopy. The advances provided by the sciDISCO protocol have the potential to rapidly advance our understanding of the cellular mechanisms of normal spinal cord function and dysfunction caused by injury or disease, thereby leading to the discovery of new therapies to improve long-term outcomes for afflicted individuals.

## Author Contributions

AB, FJ, and DM contributed to the conception and design of the study. AB, FJ, MH, and JK contributed to experimental design, data collection, and data analysis. FJ, AB, MH, and DM contributed to manuscript preparation and writing.

## Acknowledgements

We would like to acknowledge the Texas A&M University Microscopy and Imaging Center for the use of the lightsheet microscope as well as the Imaris software, particularly Dr. Holly Gibbs for her expertise and knowledge of lightsheet microscopy.

